# Molecular Characterization of Parabrachial Neurons in *Xenopus laevis* and *Silurana tropicalis*: Evolutionary Conservation and Sex-Specific Differences

**DOI:** 10.1101/2025.10.22.684011

**Authors:** Adara DeNiro, Shrinivasan Raghuraman, Kevin Chase, Baldomero M. Olivera, Ayako Yamaguchi

**Affiliations:** School of Biological Sciences, University of Utah, 257 South 1400 East, Salt Lake City, UT, 84112, USA

**Keywords:** *Xenopus*, vocalizations, parabrachial nucleus, constellation pharmacology

## Abstract

Clawed frogs communicate acoustically to coordinate reproduction, with males producing species-specific advertisement calls to attract females. In *Xenopus laevis*, males generate fast trills composed of clicks repeated at 60 Hz, a feature absent in both *Silurana tropicalis* males and *X. laevis* females, whose calls consist of slower click rates (30 Hz and 7 Hz, respectively). In male *X. laevis*, fast trills are generated by premotor neurons in the parabrachial nucleus (PBN), known as Fast Trill Neurons (FTNs). We hypothesized that FTNs are unique in male *X. laevis*, and either absent or molecularly distinct in clawed frogs that do not produce fast trills. To test this, we used constellation pharmacology to profile receptor expression of neurons via intracellular Ca²⁺ responses to pharmacological agents in PBN neurons from male *X. laevis*, male *S. tropicalis*, and female *X. laevis*. Surprisingly, we found putative FTNs in all three groups, including those that do not produce fast trills. Furthermore, a similar proportion of FTNs across groups expressed fast-kinetic voltage-gated potassium channels known to support rapid firing, indicating that the presence of these channels does not correlate with the ability to produce fast trills. Instead, some of these channels were more prevalent in males of both species compared to female *X. laevis*, suggesting a potential sex-specific, non-vocal function. The discovery of FTNs with similar molecular profiles in non-fast-trilling individuals suggests that these neurons are conserved across species and sexes, and may serve other functions. In male *X. laevis*, FTNs may have been repurposed for fast trill production during speciation. These findings provide new insight into understanding how neural circuits evolve and diversify across species and sexes.

**Summary statement:** Premotor vocal neurons share molecular profiles across clawed frogs, despite differences in calls, revealing unexpected conservation and functional divergence of homologous neurons underlying evolution of vocal behavior.

## Introduction

A primary goal of neuroscience is to understand how neural circuits coordinate behavior. Many rhythmic behaviors are generated by central pattern generators (CPG), networks of neurons that produce rhythmic motor programs in the absence of rhythmic descending input or sensory feedback (Kiehn, 2006; Marder and Bucher, 2001). A widely used approach to studying CPGs involves reduced preparations that produce fictive motor programs *in vitro*, patterns of neural activity that would generate movement if connected to muscles. These preparations allow direct electrophysiological, surgical, and pharmacological analyses of neural activity, making them invaluable tools for understanding the neural basis of behavior (Syed et al., 1990; Tarasiuk and Sica., 1997). Studies using such preparations (e.g., locomotion in lampreys, Tritonia, tadpoles, and stomatogastric system in crabs), have provided fundamental insights into the neural basis of rhythmic behavior in both invertebrates and vertebrates (Grillner, 2006; Calabrese and Marder, 2025).

Comparison of neural networks underlying homologous behaviors across species and sexes has shed light on the conserved and divergent features of neural circuit organization (Roberts et al., 2022; Katz, 2016). For example, despite exhibiting distinct feeding behaviors, the nematode species *Caenorhabditis elegans* and *Pristionchus pacificus* share homologous pharyngeal neurons and muscles (Bumbarger et al., 2013). Similarly, *Drosophila melanogaster* and *Drosophila yakuba* use homologous descending neurons to initiate species-specific courtship song in a similar social context (Ding et al., 2019). These studies show that homologous neurons can diverge to generate distinct behaviors in different species.

Previously, we showed that the calls of *Xenopus laevis* and closely related species are generated by CPGs, as isolated brains produce fictive vocalizations (Rhodes et al., 2007; Yamaguchi and Peltier, 2023). To attract females, male *X. laevis* produce advertisement calls containing two vocal phases: fast and slow trills, during which clicks are repeated at 70 Hz and 30 Hz, respectively (Figure 1A). In contrast, male *Silurana tropicalis* advertisement calls consist exclusively of slow trills (30 Hz, Figure 1B). Female *X. laevis* produce release calls composed of clicks repeated at ∼6 Hz when clasped by a male while not gravid (Figure 1C, Tobias et al., 1998). Comparison of the CPGs in these three groups of animals will provide an opportunity to understand the neural basis of vocal rhythm generation.

**Figure 1.**
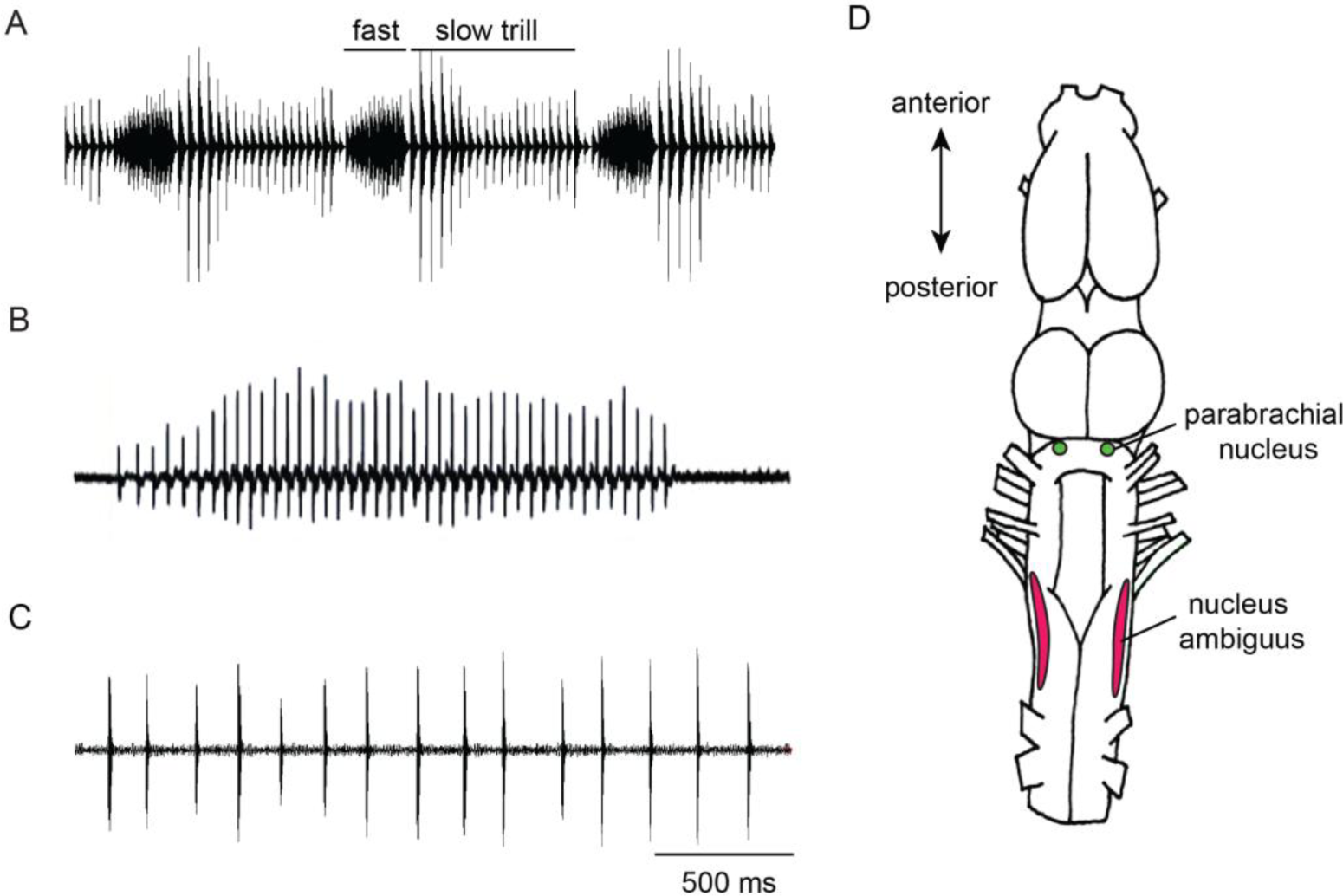
Vocalizations and central vocal pathways in Xenopus. Advertisement calls recorded from (A) male X. laevis and (B) male S. tropicalis, and a release call recorded from (C) female X. laevis. D) Schematic dorsal view of the isolated adult Xenopus brain showing the locations of key vocal nuclei, the parabrachial nucleus and the nucleus ambiguus.

In male *X. laevis*, the fast trills are generated by CPGs spanning two vocal nuclei: the parabrachial nucleus (PBN, previously referred to as DTAM) and the nucleus ambiguus (NA, laryngeal motor nucleus) (Figure 1D, Yamaguchi and Peltier, 2023; Yamaguchi et al., 2017; Zornik and Kelley, 2007). In contrast, slow trills are controlled by CPGs located in the caudal brainstem (Yamaguchi and Peltier, 2023). Interestingly, this organizational distinction of fast and slow trill CPGs is conserved across species and sexes: calls with clicks repeated at <30 Hz (e.g., advertisement calls of male *Silurana tropicalis* and the release call of female *X. laevis*) are produced by the slow trill CPGs contained in the caudal brainstem, whereas calls containing clicks repeated at >50 Hz (e.g., advertisement calls of male *Xenopus amieti*) are produced by the fast trill CPGs that span between the PBN and NA (Yamaguchi and Peltier, 2023). The lack of fast trills in male *S. tropicalis* and female *X. laevis* suggests the fast trill CPGs may be functionally absent in these animals.

A key component of fast trill CPGs in male *X. laevis* is a population of premotor neurons in the PBN, known as fast trill neurons (FTNs) (Zornik and Yamaguchi, 2012). These neurons express N-methyl-D-aspartate (NMDA) and Gamma-Aminobutyric acid (GABA) receptors (Lawton et al., 2017; Zornik and Yamaguchi, 2012), fire exclusively during fast trills, and encode both their duration and rate (Zornik and Yamaguchi, 2012). FTNs monosynaptically project to the laryngeal motoneurons in the NA (Zornik and Yamaguchi, 2012), which in turn contract laryngeal muscles to generate clicks (Kelley et al., 2017) during fast trills.

Functional differences between central vocal pathways of frogs with and without fast trills have been demonstrated through previous electrophysiological studies. Local field potential recordings from the PBN during fictive advertisement calls of male *X. laevis* reveal large-amplitude phasic activity at fast trill rates during fast but not slow trills. In contrast, PBN recordings from male *S. tropicalis* and female *X. laevis* during fictive advertisement and release calls showed little activity (Yamaguchi and Peltier, 2023). Furthermore, synapses between PBN projection neurons and laryngeal motoneurons are weaker in male *S. tropicalis* and female *X. laevis* compared to male *X. laevis* (Yamaguchi and Peltier, 2023). These findings suggest that the PBN plays a diminished role in vocal motor control in animals lacking fast trills.

Given these vocal differences, we hypothesized that molecular profiles of the PBN neurons in male *S. tropicalis* and female *X. laevis* would differ from those of male *X. laevis*. A previous study showed that female *X. laevis* PBNs contain neurons expressing NMDA receptors (NMDAR) and GABA_A_ receptors (GABA_A_R), similar to FTNs in male *X. laevis* (Inagaki et al., 2020), but their detailed molecular signature remained unclear. In this study, we compared the molecular profiles of PBN neurons in male *S. tropicalis*, female *X. laevis*, and male *X. laevis* using constellation pharmacology, a technique to identify ligand receptors and ion channels expressed by distinct neuron types through calcium imaging of dissociated neurons (Teichert et al., 2012). We specifically examined the expression of fast voltage-gated K^+^ channels (VGKCs), which support high-frequency spiking (Lovell et al., 2013; Kaczmarek, 2023; Brown and Kaczmarek, 2011), and predicted that these channels would be enriched in male *X. laevis* FTNs but absent in the other two groups.

Surprisingly, presumed FTNs that express NMDAR and GABA_A_R were found in all three groups. Further, the expression patterns of fast VGKCs among presumed FTNs were largely consistent across groups, although certain channels were more prevalent in males than in females. These findings suggest that vocal differences between species and sexes are not explained by the presence or absence of specific FTNs with fast VGKCs. Instead, the most parsimonious explanation is that FTN-like neurons with shared molecular profiles are conserved across species, serving unidentified functions, and have been evolutionarily repurposed for fast trill generation in *X. laevis* males.

## Materials and Methods

### Animals

Sixteen male *X. laevis*, seven female *X. laevis*, and seven male *S. tropicalis*, obtained from *Xenopus* 1 Corp (Dexter, MI, average +std weight = 15.91 ± 6.36 g, 20.94 ± 4.63 g, 9.21 ± 3.20 g, length = 4.77 ± 0.65 cm, 5.47 ± 0.56 cm, 3.99 ± 0.47 cm) were used for this study. Animals were housed in water-filled plastic containers within the University of Utah Animal Care Facility. Surgical procedures were performed under tricaine methanesulfonate (MS-222) anesthesia, with measures taken to minimize suffering. All the males used in the study were confirmed to be reproductively active by recording their advertisement calls before the experiment using hydrophones (Aquarian, Anacortes, WA, H2D) controlled by voice-activated recording software (Sound Analysis Pro 2011, http://soundanalysispro.com/). The sexual maturity of females was verified by the presence of fully developed ovaries at the time of surgery. This study was conducted in accordance with the National Institutes of Health’s Guide for the Care and Use of Laboratory Animals. All animal handling followed the approved protocols of the Institutional Animal Care and Use Committee (IACUC) at the University of Utah (#00001989).

### Primary Culture of Neurons

Animals were decontaminated by immersing them in tap water with 50 pg/ml gentamicin (Gibco) for a minimum of 6 hours (Laskey, 1970). Animals were fully anesthetized by injecting 0.15mg/g MS222 (1.3%, buffered to pH 7.4, filtered with 0.2um filter, Sigma-Aldrich), and decapitated. Immediately after decapitation, the isolated skull was placed on a 100mm Petri dish lined with silicone elastomer filled with ice-cold frog saline (in mM: 103 NaCl, 13 NaHCO_3_, 2 CaCl_2_, 2 KCl, 0.5 MgCl_2_, 10 HEPES and 11 dextrose, pH 7.8) oxygenated with 99% O_2_ and 1% CO_2_. The brain was then dissected out from the skull, and all meninges were removed from the brain for slice preparation.

The parabrachial nuclei (PBNs) were excised from the isolated brains using two techniques. In the first technique, 250-300μm thick transverse brain slice sections were obtained using a vibratome (Vibratome 1000) in ice-cold oxygenated saline; the PBNs were visually identified under a stereomicroscope based on the established landmark and excised using a scalpel. In the second technique, the cerebellum and optic tectum dorsal to the fourth ventricle was cut along the midline and pinned to a petri dish to expose the ventricle; the PBNs were visually identified under a stereomicroscope and excised. The first and second techniques were used on 30% and 70% of the samples, respectively. The effectiveness in selectively isolating the PBNs using the two techniques was verified by microscopic examination of the excised tissue based on distinct anatomical landmarks, such as the relative position to nuclei isthemi. Excised PBNs were transferred into HBSS (by Gibco).

The method of dissociating frog neurons was described elsewhere (Inagaki et al., 2020). Briefly, the tissue was treated with 2.5% Trypsin for 10-15 minutes, washed three times with 5 ml of Frog Neuronal Culturing Medium (FNCM, 49% Frog saline and 49% L-15 medium containing 1% Fetal bovine serum, 1% Penicillin streptomycin, with a pH of 7.8), and then triturated with a Pasteur pipet until the entire tissue was dissociated and the solution became translucent. The solution was then centrifuged at 1100 rpm for 10 minutes, and the supernatant was removed. The dissociated cell suspension (25-30μl) was applied to the center of a silicon ring (10mm ring with a 3 mm diameter hole in the center) placed on polylysine-coated culture wells (Corning). After an hour, the culturing chamber was filled with 1 ml of FNCM. Neurons were then incubated overnight for a minimum period of 12 hours (18 to 20°C for *X. laevis*, 22-26°C for *S. tropicalis* neurons).

### Calcium Imaging/Constellation Pharmacology Assay

Dissociated cells were first loaded with a FURA-2-AM dye (2.5 μM, 700 μl, 380 nm excitation/510 nm emission) for a minimum of 1 hour prior to imaging. During imaging, cells were excited with 340 nm and 380 nm light every 2 seconds, and a video image was recorded. The resulting fluorescence intensity ratio (340/380 nm) was used as an indicator of intracellular calcium concentration ([Ca²⁺]_i_) for each cell over time. Pharmacological agents (700ul-750ul, concentrations shown in Table 1) were applied manually for 10 seconds, followed by 3-4 washes using a vacuum pump controlled by a foot pedal. Recordings were obtained from 1–2 wells per animal, allowing simultaneous monitoring of hundreds of cells per well under a 10X objective. The NIS-Elements Software (Nikon Instruments, Melville, NY) was used to define regions of interest (ROIs) corresponding to each cell in a field of view and to capture the ratiometric values for each cell at each time point. The data from each ROI is represented as a time series of intercellular calcium values at 2-second intervals over the entire time-course of the experiment. This approach allows for normalization of uneven cell loading and provides measurements of [Ca²⁺]_i_ changes in response to ligand applications.

**Table 1.** Abbreviation and concentration of pharmacological agents used for constellation pharmacology

| Abbreviation | Pharmacology Agent | Concentration | Target Receptor |
| --- | --- | --- | --- |
| NMDA | N-methyl-d-aspartate | 200 μM | NMDA-receptor |
| Sub P | Substance P | 1 μM | Neurokinin-1-receptor |
| ACh | Acetylcholine | 1 mM | Acetylcholine-receptor |
| GABA | γ-Aminobutyric acid | 1 mM | GABAergic receptor |
| Gly | Glycine | 1 mM | Glycinergic-receptor |
| d-Ser | d-Serine | 20 μM | NMDA-receptor |
| BDS-I | N/A | 5 nM | K<sub>v</sub>3.1, 3.2, 3.4 |
| κM-MuIIIA | N/A | 1 μM | K<sub>v</sub>1.1 and 1.2 |
| TEA | Tetraethylammonium | 10mM | Nonspecific K<sub>v</sub> channels |
| 4-AP | 4-aminopyridine | 1mM | Nonspecific K<sub>v</sub> channels |

An increase in cytosolic Ca^2+^ concentration induced by the application of high-concentration KCl (15mM or 30mM), which depolarizes the membrane potential due to the shift in the K⁺ equilibrium potential, was used to distinguish neurons from glial cells, as voltage-gated Ca²⁺ channels are present in neurons but absent in glial cells (Carmignoto et al., 1998; Teichert et al., 2012).

We next applied a panel of pharmacological agents sequentially to ∼100 to 500 dissociated neurons and monitored intracellular calcium concentration ([Ca²⁺]_i_). The activation of receptors and ion channels that depolarized the membrane potential was measured as an elevated [Ca²⁺]_i_ (Figure 2). The hyperpolarized membrane potential (i.e., GABA in this study) was detected using the following method. Three pulses of 15mM KCl were applied at 5 to 7-minute intervals, while the cells were incubated with inhibitory ligands between the first and second pulses. A reduced amplitude of [Ca²⁺]_i_ after GABA incubation by the second KCl pulse compared to the first pulse was taken as evidence of reduced depolarization, due to an increased influx of Cl-mediated by the GABA_A_ receptors (Figure 2A, B). Recovery of the [Ca²⁺]_i_ signal in response to a third KCl pulse confirmed that the reduced depolarization was due to the activation of GABA_A_ receptors, not due to deteriorated cell health.

**Figure 2.**
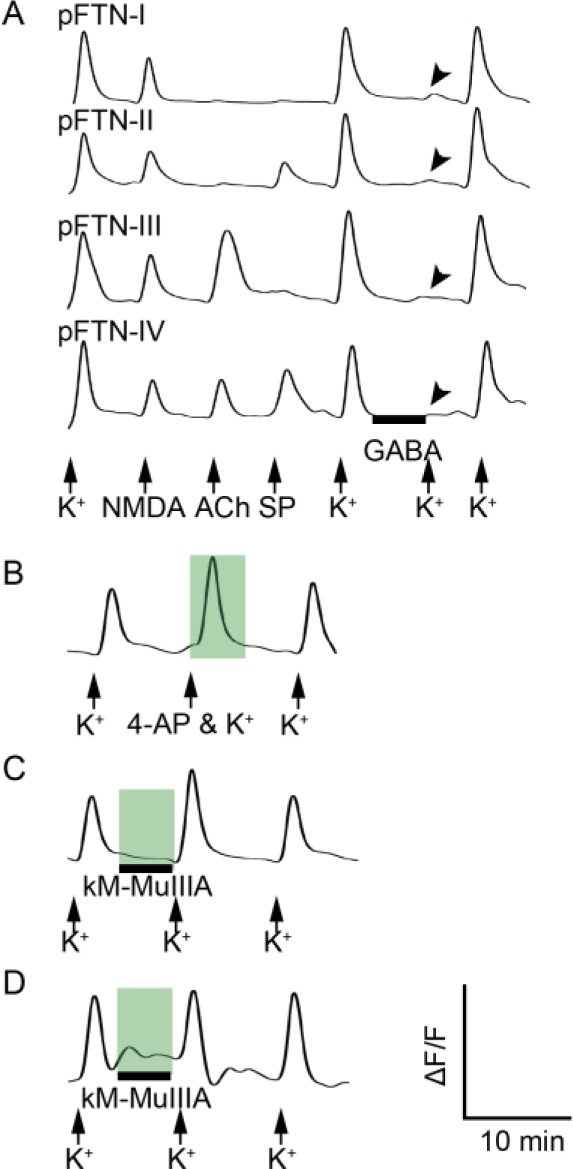
Example calcium imaging traces from parabrachial neurons. The y-axis represents relative cytoplasmic Ca^2+^ concentration ([Ca^2+^]_i_), and the x-axis represents time. Pharmacological agents were applied at time points indicated by black arrows. Solid black bars denote periods of incubation with the respective pharmacological agents. A: Example traces from pFTN subtypes I through IV. Three pulses of K^+^ were applied at the end of each trace. Between the first and the second pulse, neurons were incubated with GABA. The absence of [Ca^2+^]_i_ signals in response to the second pulse (indicated by gray arrowheads), which recovers by the third pulse, is interpreted as evidence for GABA_A_ receptor expression. pFTN-I respond to NMDA and GABA, but not to ACh or SP. pFTN-II respond to NMDA, GABA, and SP, but not to ACh. pFTN-III respond to NMDA, GABA, and ACh, but not to SP. pFTN-IV respond to NMDA, GABA, SP, and ACh. B. Amplification of the KCl-induced Ca^2+^ signal (highlighted with green background) when co-applied with 4-AP. C. Amplification of the KCl-induced Ca^2+^ signal (green background) following incubation with κM-MuIIIA. D. Direct increase in [Ca^2+^]_i_ (green background) in response to κM-MuIIIA application, presumably due to reduced K^+^ currents at the resting membrane potential.

To determine the presence of fast voltage-gated K^+^ channels (VGKCs), we used two approaches. In both approaches, three pulses of 15mM KCl were applied, and the response to the third pulse was used to confirm the health of the cells, as described above. In the first approach, the second KCl pulse contained an antagonist for fast VGKCs. An increase in the amplitude of [Ca²⁺]_i_ in response to the second pulse compared to the first pulse is used as evidence for the presence of the target K^+^ channels, as the absence of voltage-gated K^+^ conductance should result in an increased membrane potential depolarization (Figure 2C). In the second approach, the cells were incubated with an antagonist between the first and second pulses. An increase in [Ca²⁺]_i_ of the second pulse was taken as evidence for the presence of target K^+^ channels (Figure 2D), and an increase in [Ca²⁺]_i_ during the incubation with an antagonist indicated the presence of the target channel active at the resting membrane potential (Figure 2E). (Yanagihara and Giglio, 2024).

### Data analyses

A single calcium imaging experiment measured the intracellular calcium concentration for 100s of cells at 2 s intervals over a time course of up to 1.5 h. The time course of the [Ca²⁺]_i_ trace (i.e., 340:380 ratio) for each cell (ROI) trace was analyzed using a set of R functions (R Core Team, 2021). Traces with abnormal response patterns were removed before the analysis. The MALDIquant package (Gibb and Strimmer, 2012) was used to correct baselines (estimateBaseline function with the “SNIP” method), smooth traces (smoothIntensity function with the “SavitzkyGolay” method with a halfWindowSize of 3), and detect peaks (detectPeaks function with the “MAD” method with a halfWindowSize of 30 and an SNR.lim setting of 4). Peak heights were calculated by subtracting the baseline from the maximum deflection of the 340:380 ratio within a defined window for each agonist. Traces were further processed using a second smoothing function (smooth.spline with a smoothing parameter (spar) of 0.3) and normalized by subtracting its minimum value and scaling it to a range of 0 to 1, facilitating comparison across traces.

We applied a signal-to-noise (SNR) threshold of 4 times the standard deviation of the noise amplitude, with a minimum value of 0.05 above the [Ca²⁺]_i_ baseline to minimize false positives in regions with low variance in the 340:380-nm ratio. All ROIs were scored as a yes/no response to each ligand application. Subthreshold signals were labeled as “no” response. The criteria gave a false-positive rate <0.001 (e.g., peaks during times of no input). A “yes” score was given if 1) the application of an excitatory agonist elicited a peak in the response window of 30 s, 2) the incubation with an VGKC antagonist alone elicits an increase in [Ca²⁺] _i_, (i.e., direct effect), 3) the incubation of an VGKC antagonist is followed by the amplified KCl-induced [Ca²⁺]_i_ relative to the baseline signal of the KCl-induced [Ca²⁺] before antagonist application (i.e., indirect effect), 3) the incubation with an VGKC antagonist alone elicits an increase in [Ca²⁺]_i_, (i.e., direct effect), 4) the co-application of an VGKC antagonist with KCl amplified the [Ca²⁺]_i_ response, or 5) the incubation with an inhibitory ligand reduced or completely eliminated the following KCl-induced [Ca²⁺]_i_ signal. For the final analyses, all cells that exhibited direct or indirect effects were grouped together.

In addition to yes/no response, we estimated the total [Ca²⁺]_i_ over time by quantifying the area under the curve (AUC) of the responses. The effects of the K^+^ channel antagonists were determined by calculating the AUC difference between the KCl-induced response after incubation (a) and before incubation (b) divided by (a) to calculate the percent increase or decrease. To normalize this effect, a relative ratio was employed using the formula (b-a)/(b+a), allowing for a standardized comparison across conditions. AUC relative ratios ranged from 0 to 1 for amplified responses and 0 to -1 for inhibited responses.

### Statistical Analyses

We created a generalized linear mixed-effects model (GLMM) using the “glmer” function from the R package lme4 (Bates et al., 2015) to test the effects of neuron type, species or sex groups (species and sex), and their interaction. The model included “Neuron Type” (the classes of putative fast trill neurons) and “Species/Sex Group” as fixed effects, with “Well” (e.g., wells that contained dissociated cells) treated as a random effect to account for non-independence within wells. For binomial outcomes (yes/no response), we specified *Y_ij ∼ Bernoulli(p_ij)*, *logit(p_ij) = η_ij = X_ij β + b_j, p_ij = 1 / (1 + exp(−η_ij))*, where *i* index observations (cells), *j* index wells, *Xij* represent the fixed-effects design row (intercept, Neuron Type, Species/Sex Group) and *β* the fixed-effects coefficient vector. The random intercept for Well *j* is *b_j*, modeled as *b_j* ∼ *N(0, σ_b#x005E;2)*. The GLMM estimated the probability of a successful outcome for each observation by transforming the linear predictor (log-odds) into probabilities using the inverse logit function. This approach provided interpretable probability estimates ranging from 0 to 1. For continuous outcomes (e.g., AUC relative ratios), we used a Gaussian mixed model: *Y_ij = X_ij β + b_j + ε_ij,* with *ε_ij ∼ N(0, σ_ε#x005E;2)*. Standard errors and 95% Wald confidence intervals were computed as β#x005E; ± 1.96 SE(β#x005E;), and significant difference among fixed effects was assessed with Wald z tests (z = β#x005E;/SE). Statistical analyses and data processing were conducted in RStudio (R Core Team, 2021).

## Results

### The presence and proportion of putative fast trill neurons in the parabrachial nucleus do not correlate with fast trills in the vocal repertoire

Although male *S. tropicalis* do not produce fast trills, we found putative FTNs (pFTNs) in their PBN, as determined by their responsiveness to NMDA with d-serine and GABA (Figure 3). We also confirmed previous findings that female *X. laevis* PBNs also contain pFTNs despite a lack of fast trills (Inagaki et al., 2020). In male *X. laevis*, the proportion of pFTNs and non-pFTNs is nearly equal (Figure 3A). However, the proportion of pFTNs was significantly higher than non pFTNs in both *S. tropicalis* and female *X. laevis* (GLMM, *S. tropicalis* male: β= 0.79718 ± 0.12475, z= 6.390, p<0.001; *X. laevis female*: β= 0.99600 ± 0.12972, z= 7.678, p<0.001; Figure 3A). The proportion of pFTNs in *S. tropicalis* and female *X. laevis* was significantly higher, while the proportion of non-pFTNs of these groups was significantly lower, compared to male *X. laevis* (GLMM: *S. tropicalis* male vs. *X. laevis* male: β= 0.69372 ± 0.07, z= 9.896, p<0.001; *X. laevis* female vs. *X. laevis* male: β= 0.89253 ± 0.0786, z= 11.362, p<0.001; Figure 3A). Thus, contrary to our expectation, pFTNs are more prevalent in frogs that do not produce fast trills than in those that do.

**Figure 3.**
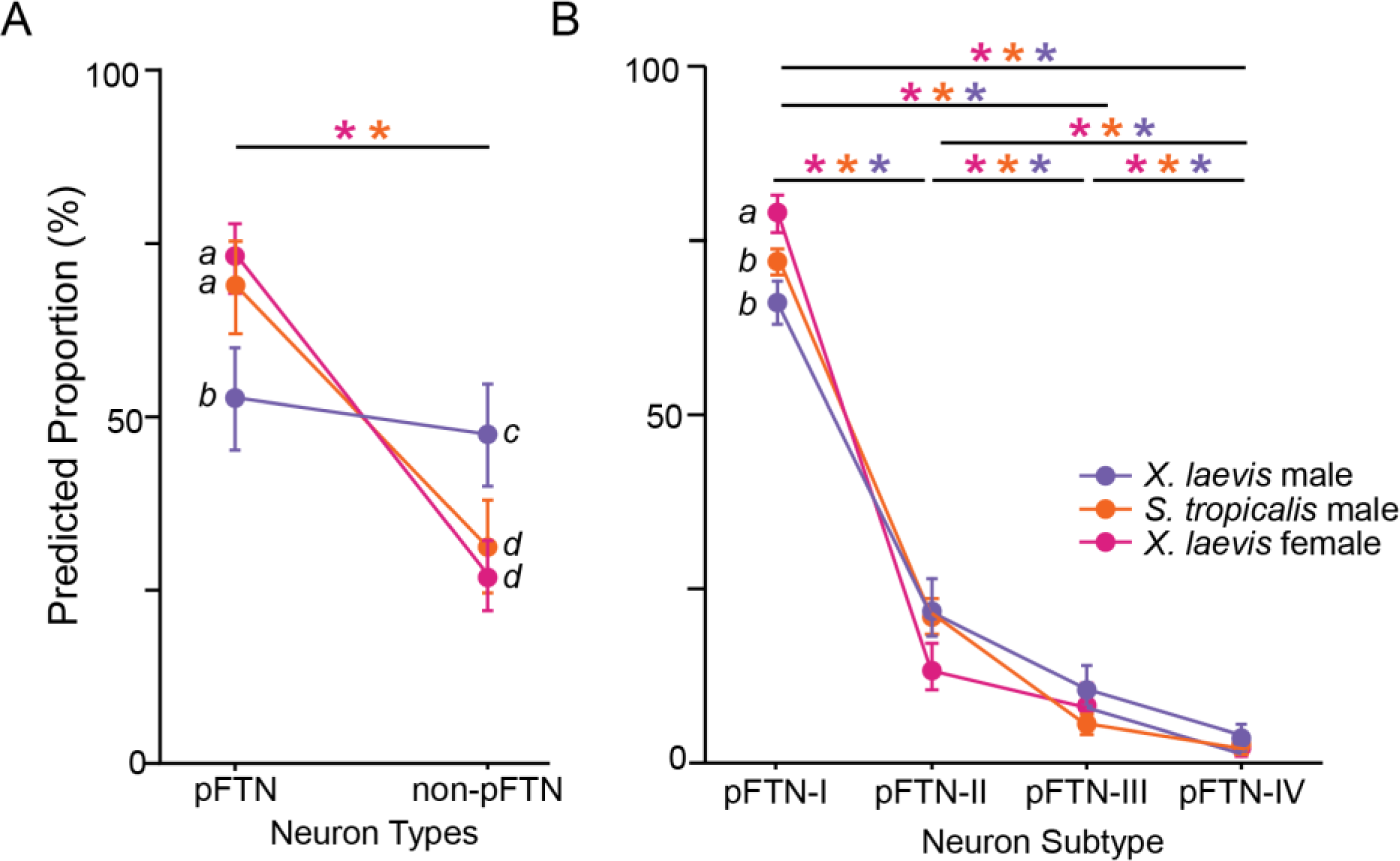
Proportion of pFTNs and non-pFTNs. A. Line graph of the predicted proportion of pFTNs and non-pFTNs of X. laevis male (n=15), X. laevis female (n=7), and S. tropicalis male (n=7). Each colored circle represents the predicted proportion, with the error bars indicating the 95% confidence intervals. Significant differences across pFTNs and non-pFTNs within each group are marked by asterisks over solid bars. Difference across species/sex groups within each neuron type are denoted by letters: pFTN groups labeled “a” and “b” are significantly different, as are non-pFTN groups labeled “c” and “d”. B. Line graph showing the predicted proportion of four pFTN subtypes in the same three groups. Circles represent predicted proportions, with error bars indicating 95% confidence intervals. Significant differences across neuron subtypes within each group are indicated by an asterisk over a solid bar; differences across species/sex groups within each subtype are denoted by letters.

Previously, we demonstrated that pFTNs of male and female *X. laevis* can be classified into four subgroups based on their responsiveness to acetylcholine and substance P, with subtypes I, II, III, and IV, ranked from most to least common in both sexes (Inagaki et al., 2020). In the present study, we found that male *S. tropicalis* also possesses all four pFTN subtypes (Figure 2, 3B) with proportions similar to those of male and female *X. laevis* (Figure 3B). Notably, the proportion of subtype pFTN-I in female *X. laevis* was significantly higher than in both male groups (GLMM: *X. laevis* female vs. *S. tropicalis* male: β= 0.386 ± 0.11, z= 3.528, p<0.001; *X. laevis* female vs. *X. laevis* male: β= 0.66017 ± 0.09, z= 6.90, p<0.001; Figure 3B). Taken together, these results suggest that the presence and proportion of pFTNs and their subtypes do not correlate with the presence of fast trills in the vocal repertoire. Instead, the data suggest that subtype pFTN-I may be more abundant in females than in males, regardless of species.

### Fast-kinetic voltage-gated K^+^ channel expression in pFTNs does not correlate with the presence of fast trills in their vocal repertoire

Given that all three species/sex groups have pFTNs in the PBNs despite their differences in fast trills production, and that the two groups without fast trills exhibit a higher proportion of pFTNs, we hypothesized that pFTNs of female *X. laevis* and male *S. tropicalis*, defined by the presence of two receptor types (GABA_A_ and NMDAR), may differ functionally from those in male *X. laevis*. Specifically, we reasoned that fast kinetic voltage-gated K+ channels (VGKC), known to control rapid and precisely timed spiking (Lovell et al., 2013; Kaczmarek, 2023; Brown and Kaczmarek, 2011), might distinguish pFTNs of male *X. laevis* from those of male *S. tropicalis* and female *X. laevis*. To this end, we used antagonists for fast VGKCs, 4-AP, κM-MuIIIA, and BDS-I, to determine whether these channels are more prevalent amongst male *X. laevis* pFTNs than in male *S. tropicalis* and female *X. laevis*.

The proportion of pFTNs responsive to 4-AP, a broad-spectrum fast VGKC antagonist, was significantly higher in male *X. laevis* than in male *S. tropicalis* (GLMM: *S. tropicalis* male vs. *X. laevis* male: β= 0.34852 ± 0.0916, z= 3.803, p<0.001; Figure 4A), but the proportion of 4-AP sensitive pFTNs in females did not differ from either male groups. Further, the proportion of 4- AP-sensitive neurons was significantly higher in pFTNs than in non-pFTNs in both male *X. laevis* and male *S. tropicalis* (GLMM: *S. tropicalis* male pFTN vs. non-pFTN: β= -0.661 ± 0.153, z= - 4.319, p<0.001; *X. laevis* male pFTN vs. non-pFTN: β= -0.103 ± 0.09, z= -10.856, p<0.001; Figure 4A), but not in female *X. laevis*. At the subtype level, the proportion of 4-AP sensitive pFTN-I was significantly higher in male *X. laevis* than in male *S. tropicalis*, but that of female *X. laevis* did not differ from either males (GLMM: *S. tropicalis* male vs. *X. laevis* male: β= 0.389 ± 0.106, z= 3.462, p<0.001; Figure 4B). The results indicate that the proportion of 4-AP-sensitive pFTN neurons does not distinguish fast trillers from slow trillers, but may reflect species-specific differences between the male groups.

**Figure 4.**
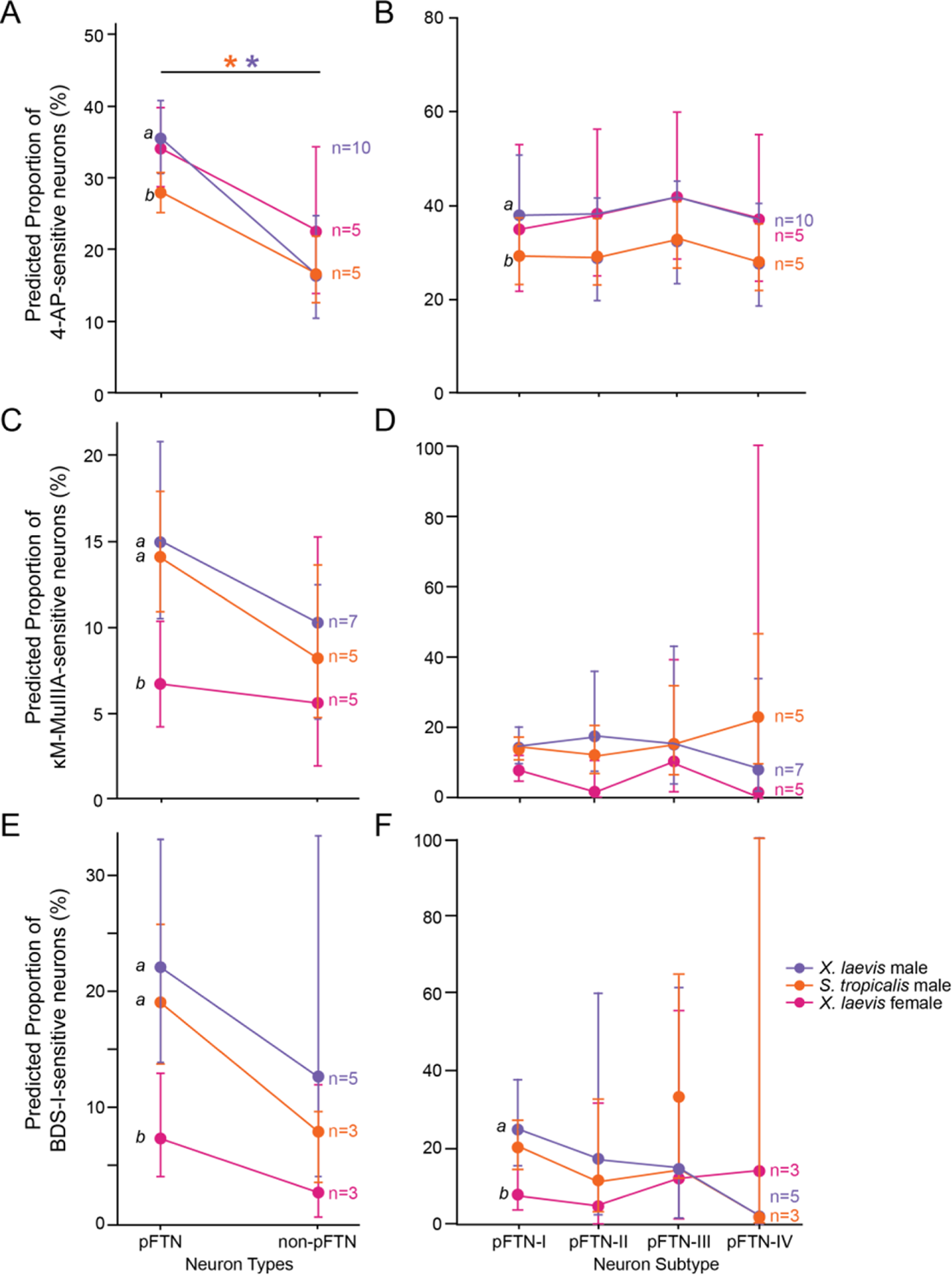
Predicted proportion of voltage-gated K^+^ channel sensitive neurons in pFTNs and non-pFTNs. A, C, and E: Line graph showing the predicted proportion of 4-AP- (A), κM-MuIIIA- (C), and BDS-I- (E) sensitive cells among pFTNs and non-pFTNs of X. laevis males, X. laevis females, and S. tropicalis males. Each circle represents the predicted proportion, with error bars indicating 95% confidence intervals. Significant differences across neuron type within each group are indicated by asterisks and solid bars; differences across species/sex groups within each neuron type are denoted by letters. B, D, and F: Line graph showing the predicted proportion of 4-AP- (B), κM-MuIIIA- (D), and BDS-I- (F) sensitive cells across four pFTN subtypes in the same three groups. Each data point represents the predicted proportion, with error bars indicating upper and lower 95% confidence intervals. Each circle represents the predicted proportion, with error bars indicating 95% confidence intervals. Significant differences across neuron subtype within each group are indicated by asterisks and solid bars; differences across species/sex groups within each neuron subtype are denoted by letters.

Since 4-AP targets multiple Kv channels, we next investigated if specific types of VGKCs differ across species/sex groups using selective antagonists for Kv1.1 and 1.2 (κM-MuIIIA), and for Kv3.1, 3.2, and 3.4 (BDS-I, Yeung et al., 2005; Ciccone et al., 2019). The results showed nearly equal proportions of BDS-I- and κM-MuIIIA-positive neurons were pFTN and non-pFTNs in all sex/species group, indicating that these channels are not exclusive to pFTNs. However, when comparing across species/sex groups, the proportions of BDS-I and κM-MuIIIA sensitive pFTNs were significantly higher in both male groups than in female *X. laevis,* (GLMM: *X. laevis* female vs. *S. tropicalis* male (κM-MuIIIA): β= 0.831 ± 0.198, z= 4.205, p<0.001; *X. laevis* female vs. *X. laevis* male (κM-MuIIIA): β= 0.903 ± 0.186, z= 4.85, p<0.001; *X. laevis* female vs. *S. tropicalis* male (BDS-I): β= 1.095 ± 0.256, z= 4.28, p<0.001; *X. laevis* female vs. *X. laevis* male (BDS-I): β= 1.278 ± 0.223, z= 5.74, p<0.001; Figure 4C, E), while non-pFTNs showed no significant differences across groups. These results suggest a sex-specific, and not vocal-related difference, in fast VGKC expression within pFTNs.

Further analysis of pFTNs subtypes to the antagonists revealed that the proportion of BDS-I-sensitive pFTN-I of male *X. laevis* was significantly higher than that of female *X. laevis* (GLMM: *X. laevis* female vs. *X. laevis* male: β= 1.383 ± 0.253, z= 5.46, p<0.001, Figure 4F), while all other subtypes showed similar proportions across groups. These results indicate that none of these VGKC types reliably distinguishes pFTNs of fast trillers from those of slow trillers. Instead, the results suggest that the proportion of pFTNs with κM-MuIIIA- and BDS-I-sensitive K^+^ channels reflects sex differences rather than vocal behavior.

Although the proportion of neurons with fast kinetic VGKC in pFTNs did not correlate with fast trills, we further explored whether the VGKC conductance in pFTNs might account for the vocal differences. If so, such differences should be reflected in the relative magnitude of KCl-induced intracellular calcium ([Ca²⁺]_i_) in the presence of fast VGKC antagonists. To this end, we quantified the relative magnitude of calcium signals by comparing the ratio of the area under the curve (AUC) with and without antagonists (see Methods). Cells were incubated with κM-MuIIIA and BDS-I, whereas 4-AP was co-applied with KCl since incubation with 4-AP appear to deteriorate the health of the neurons. Our results showed that the predicted AUC relative ratios for 4-AP, κM-MuIIIA, and BDS-I were similar between pFTNs and non-pFTNs (Figure 5A, 5C, 5E), and across all pFTN subtypes and species/sex groups (Figures 5B, D, F), indicating that the pFTNs are not only similar in proportion, but also in VGKC conductance across all three groups.

**Figure 5.**
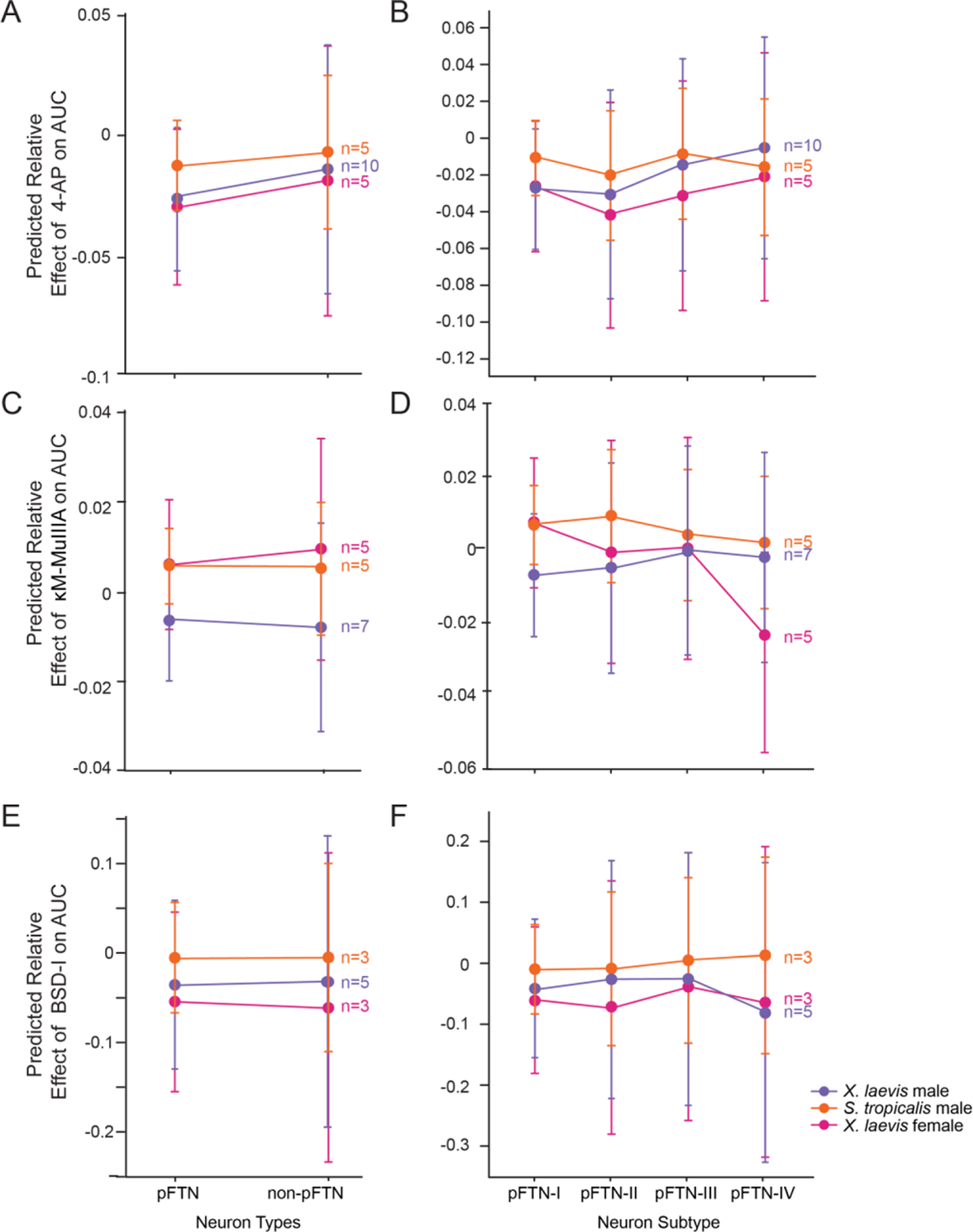
Predicted relative effect of VGKC antagonists on the area under the curve (AUC) of [Ca^2+^]_i_ in pFTNs and non-pFTNs. A, C, and E: Line graph showing the predicted relative effect of 4-AP (A), κM-MuIIIA (C), and BDS-I (E) on the AUC among pFTNs and non-pFTNs of X. laevis males, X. laevis females, and S. tropicalis males. B, D, and F: Predicted relative effect of 4-AP (B), κM-MuIIIA (D), and BDS-I (F) on the AUC across four pFTN subtypes of X. laevis males, X. laevis females, and S. tropicalis males. Each circle represents the predicted probability, with error bars indicating 95% confidence intervals.

In summary, we found no systematic difference in the receptors and ion channels expressed in the pFTNs across species and sexes, despite differences in vocal behavior. Instead, our data reveals a sex-specific pattern in the proportion of pFTNs expressing fast VGKC.

## Discussion

The PBN of male *X. laevis* exhibits large phasic activity during fictive fast trills but not during slow trills (Zornik and Yamaguchi, 2012; Yamaguchi and Peltier, 2023). In contrast, the PBNs of *S. tropicalis* and female *X. laevis* show minimal activity during fictive calls (Yamaguchi and Peltier, 2023), suggesting the involvement of PBNs only during fast trills regardless of the sex and species. This study aimed to determine whether parabrachial neurons in three groups of frogs, those with and without fast trills, exhibit distinct expression patterns of receptors and ion channels. Although male *X. laevis* are the only group with fast trills, and their FTNs generate them, we identified putative FTNs in all three groups. Surprisingly, the proportion of pFTNs was significantly higher in slow trillers (male *S. tropicalis* and female *X. laevis*) than in male *X. laevis*. Additionally, female *X. laevis* had a significantly higher proportion of pFTN-I neurons compared to males in either species. Similarly, the expression patterns of fast-kinetic voltage-gated K^+^ channels did not align with the presence of fast trills in the vocal repertoire, but correlate more strongly with sex. We suggest three possible explanations for the results.

First, pFTNs in the three groups of animals may differ in their molecular profiles. In our study, we used seven ligands for channels and receptors suspected to be involved in fast trill generation to classify neurons into pFTNs. However, our preliminary RNA sequencing data show that brainstem neurons of male and female *X. laevis* express more than 40,000 types of transcripts. Similarly, the calcium signal measurements used in this study may not precisely capture changes in spike frequency, potentially overlooking subtle ligand-induced changes in neuronal activity. Thus, single cell RNA sequencing and electrophysiological recordings of the pFTNs may reveal systematic differences between fast and slow trillers.

Second, the molecular profiles and the intrinsic properties of pFTNs in fast and slow trillers may be identical, yet their functions may have diverged between the two groups. It remains unclear which vocal CPGs, fast or slow, is ancestral in the genus *Xenopus*, as both fast and slow trillers are widely distributed across clades (Tobias et al., 2011; Evans et al., 2015). One possibility is that pFTNs may serve a universal function in all *Xenopus* species but were co-opted by fast trillers to participate in fast trill CPGs. Alternatively, pFTNs may have evolved specifically for fast trill generation, with slow trillers repurposing them for other functions. Notably, the synapses between PBN projection neurons and laryngeal motoneurons are weaker and less reliable in male *S. tropicalis* and female *X. laevis* than in fast clickers (Yamaguchi and Peltier, 2023), suggesting that differences in synaptic connectivity, rather than changes in intrinsic properties, may explain the functional divergence between the two groups.

Third, the pFTNs found in slow trillers may be an evolutionary vestige and do not partake in generating fast trill. Similar mismatches between neurons, neural circuitry, and behavior are evident in other species. For example, flight neural networks homologous to those of locusts are found in the flightless grasshopper (Arbas, 1983), and fully functional song motor pathways, which can artificially be activated, are present in female *Drosophila melanogaster* that never express the behavior (Clyne et al., 2008). Thus, pFTNs may have evolved to generate fast trills and are inherited by all *Xenopus* species regardless of the presence or absence of fast trills, and remain latent in their CNS.

Interestingly, our results suggest sex-specific differences pFTNs. Female *X. laevis* exhibited a significantly higher proportion of pFTN-I than in either males (Figure 3B). In contrast, the proportions of κM-MuIIIA- and BDS-I-sensitive pFTNs were higher in male *X. laevis* and male *S. tropicalis* than in female *X. laevis* (Fig 4C, E), and 4-AP-sensitive neurons are more prevalent in pFTNs than in non-pFTNs in both males, but not in female *X. laevis* (Figure 4A). There are ample examples of sexually distinct expression of VGKCs. For instance, electrocytes of the electric organ of the weakly electric fish, *Sternopygus*, were found to have higher expression of Kv1.1a and Kv1.2a in females than in males, supporting the sexually distinct frequency of electric organ discharge (Few and Zakon, 2007). Similarly, the laryngeal motoneurons of *X. laevis* have sexually distinct conductance of low-threshold K^+^ currents, facilitating precise spike timing in males (Yamaguchi et al., 2003), and hypothalamic gonadotropin-releasing hormone (GnRH) neurons in mice show sexually distinct changes in voltage-gated K^+^ currents during temporal lobe epilepsy (Rajan & Christian-Hinman, 2024). The PBN is known to regulate respiration, sensory integration, and homeostatic regulation in mammals (Pauli et al., 2022), with similar functions identified in amphibians (Zornik and Kelley 2008). Although sex-specific functions of PBN neurons in *Xenopus* remain unclear beyond vocal production, it is plausible that sexually distinct sensory processing in the PBN is mediated by neurons with differential VGKC expression.

In conclusion, this study showed that the presence of putative fast trill neurons and the expression of fast kinetic VGKC do not directly predict the production of fast trills in *Xenopus*. Instead, the pFTNs in fast trillers may have acquired function to be a part of fast trill CPGs by forming novel synaptic connectivity while those in slow trillers serve different functions, or remain latent as evolutionary vestiges. By examining how pFTNs integrate into vocal CPGs to produce sex and species-specific calls, we can gain a better understanding of the role these differences play in reproductive isolation and speciation. Additionally, the observed sex-specific differences in fast-kinetic voltage-gated K+ channel sensitivity raise important questions about the role of hormonal regulation in shaping neuronal excitability.

## Data Availability

The data that support the findings of this study are available from the corresponding author upon reasonable request.

## Acknowledgments

We thank Jody Reimer and Fred Adler for their assistance with data analyses. This work was supported by NSF IOS-1934386 (AY).

## Author Contributions

A.D, S.R., B.M.O., and A.Y. conceived and designed research; A.D performed experiments; A.D and K.C. analyzed data; A.D, S.R., B.M.O., and A.Y. interpreted results of experiments; A.D and A.Y prepared Figures; A.D and A.Y. drafted manuscript; A.D, S.R., K.C., B.M.O and A.Y. edited and revised manuscript; A.D, S.R., K.C., B.M.O., and A.Y. approved final version of manuscript.

